# Ornament, armament, or toolkit? Modelling how population size drives the evolution of birdsong, a functional cultural trait

**DOI:** 10.1101/2021.04.29.442039

**Authors:** Emily J. Hudson, Nicole Creanza

## Abstract

Oscine songbirds have been an important study system for social learning, particularly because their learned songs provide an analog for human languages and music. Here we propose a different analogy; from an evolutionary perspective, could a bird’s song be more like an arrowhead than an aria? We modify a model of human tool evolution to accommodate cultural evolution of birdsong: each song learner chooses the most skilled available tutor to emulate, and each is more likely to produce an inferior copy than a superior one. Similarly to human tool evolution, we show that larger populations foster improvements in song over time, even when learners restrict their pool of tutors to a subset of individuals. We also demonstrate that songs could be simplified instead of lost after population bottlenecks if lower-quality traits are easier to imitate than higher-quality ones. We show that these processes could plausibly generate empirically observed patterns of song evolution, and we make predictions about the types of song elements most likely to be lost when populations shrink. More broadly, we aim to connect the modeling approaches used in human and non-human systems, moving toward a cohesive theoretical framework that accounts for both cognitive and demographic processes.

## Introduction

Social learning—“learning that is facilitated by observation of, or interaction with, another individual or its products” [1]—is pervasive across the animal kingdom, from invertebrates to cetaceans and primates [2–4]. Social learning is a special case of inheritance that provides non-genetic pathways for adaptive traits to be passed from one generation to the next. How does the pattern and process of evolution differ when important traits are learned rather than transmitted genetically? The discovery of genes as the primary unit of inheritance meant that Darwin’s ideas about natural selection could be synthesized with an underlying mechanism of transmission. A similar synthesis between genetics, evolutionary processes, and broad behavioral patterns is still needed for learned traits. In combination with experiments investigating the mechanisms of learning, mathematical models can help generate hypotheses and test assumptions about cognitive processes related to the transmission of social information.

The oscine songbirds offer a promising system for modeling the interaction of evolution and culture; like humans, songbirds must undergo a learning process to produce effective vocal communications. In contrast to language, however, birdsong is often shaped by the transmission properties of the environment, such as foliage density [5,6] or background noise [7]. Most notably, much research on birdsong has focused on social selection, including female preferences for song elaboration or vocal performance (reviewed in [8,9] but see [10]). These preference-shaped features are (to varying degrees) socially transmitted and constrained by the availability of tutors. For example, even if the song trait of repertoire size has genetic and developmental components [11–13], pupils with a small pool of potential tutors will be limited in the types of sounds from which to build their repertoires [14]. Returning to an analogy with human culture, then, subtle variations in a bird’s song can potentially be linked to fitness differences, much like the skill to craft a sharp arrowhead can translate to increased fitness for a human hunter. In both cases, a pupil’s learned skill depends on the social environment in which it learns.

Here, we propose that treating birdsong as a functional tool is a novel and complementary approach to analogizing birdsong to human language. Although songs, unlike tools, are not physical objects, songs have fitness consequences and may experience cumulative cultural change towards a more effective version (which rarely happens in language). By emphasizing song’s functional aspects, we gain new methods to understand its evolutionary dynamics and how demographic factors affect song over time. We use a cultural evolutionary approach that has been applied to social learning and human tool evolution [15] to make predictions about song evolution. In essence, we represent song as a difficult skill that all learners in the population attempt to reproduce with varying success. This simple but powerful framework has yielded novel insights into human behavior (e.g. [16]; discussed in more detail below), and it holds promise to do the same for cultural traits in other species. In particular, this approach provides a specific mechanism to explain why attractive elements of song are sometimes lost when bird species expand into new habitat and experience an associated population bottleneck. Unlike cultural drift, which can be dominated by random processes, the model we propose here incorporates selection—the assumption that certain forms of a cultural trait are beneficial or preferable—while integrating population demographics and specific aspects of cognitive ability [17–20]. Additionally, we apply the tools of social network analysis to show how changing population connectivity affects trait evolution. Finally, we model the outcome on songs if trait difficulty varies with the elaboration of the trait, for example, if less attractive songs are easier to learn and transmit than more attractive ones.

## Basic model

In the tradition of anthropologists and cultural evolutionary theorists [21–23], Henrich [15] applied mathematical models to show that the progressive loss of complex skills observed across several thousand years of the Tasmanian archaeological record was best explained as an outcome of demographic factors (namely, effective population size). In Henrich’s mathematical model (based on the Price equation; see Supplement for more details), a learned trait (such as a netweaving technique, or the straightness of an arrow shaft, p. 200 of ref. [15]) gradually improves or declines across generations in a culture, depending on the interaction of three factors: 1) how difficult the trait is to accurately copy (α), 2) how widely learners vary in their attempts to copy the trait (β), and 3) how many learners and potential tutors exist in the current population (*N*), a non-cognitive factor that nevertheless plays an important role in cultural change.

Given a trait that is difficult to learn, most individuals will produce inferior imitations, and thus only sufficiently large populations will contain enough pupils to consistently produce slightly improved imitations by chance, therefore increasing the maximum value of the trait in the next generation (Figure 1). Importantly, and perhaps counterintuitively, the population variation in skill of available tutors does not influence trait change in the next generation in this model; only the highest-skill tutor is chosen by pupils to attempt to imitate. This process differs from drift in that traits that decline in skill level over time are not random, but rather tend to be the most difficult traits to imitate [15]. While the application of this model to empirical evidence in human cultural evolution has been widespread, if contentious (e.g. [24–26]), to our knowledge it has never been applied to non-human cultures.

**Figure 1.**
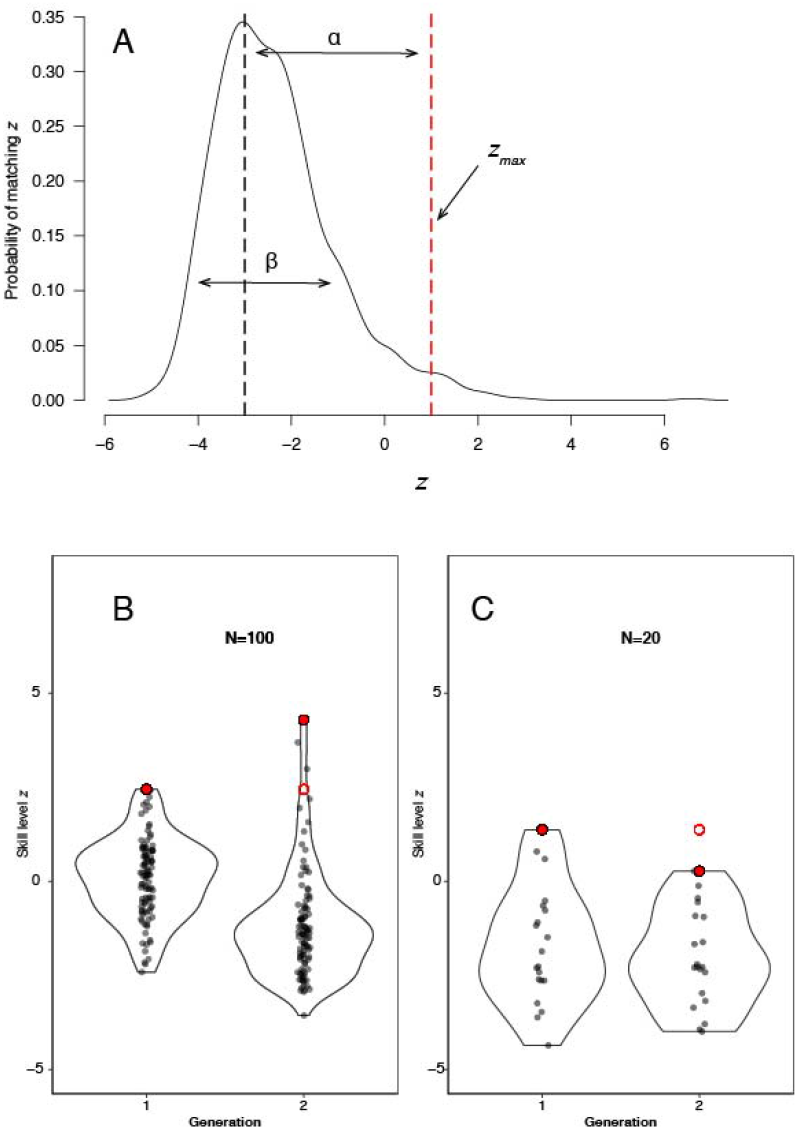
**(A)** The probability distribution of a learner attempting to copy a trait with the highest *z* value possible (*N*=100 individuals). The red line indicates the maximum *z* value of the tutor population, *z*_max_. Most pupils in the population will learn to produce the trait with *z* values below *z*_max_; by chance, however, some individuals will produce imitations above *z*_max_, such that large populations are more likely to exceed the previous maximum value in the population. **Lower panels:** The change in *z_max_* from the first to second generation when population size is 100 (**B**) and 20 (**C**). The second generation learns from the first generation, attempting to copy the most skilled (highest *z*) individual (filled red circle on the left). The open circle shows the maximum *z* value of the previous generation. Whenever the new *z_max_* exceeds that of the previous generation, the trait’s skill level will improve over time; if *z_max_* decreases, the trait will degrade (**C**).

## Applying the model to birdsong: methods and results

To apply the above model to birdsong, it is necessary to translate the concept of an individual’s skill along a continuous axis (*z*) to a measurable feature of song. Linking specific song traits to fitness is challenging, but some authors have shown how reproductive success relates to particular performance traits in many taxa, including oscine songbirds (reviewed in [27]); for example, female preference for rapidly trilled notes [28–31], large repertoire size [32–34], and the production of particular elements [35–38]. In this model, we define a better song (higher *z*) loosely as any song that increases the singer’s fitness. In most oscine birds, this concept can also be expressed as a song that is more attractive to the receiver (usually, a prospective female mate). Although not all components of birdsong are learned, for the predictions of the Price equation to apply, it is only necessary for some degree of social inheritance to take place; song traits with partial genetic control can still be modelled if the distribution of learner performance is affected by the availability of skilled tutors.

Henrich’s model assumes that, due to the incentive to produce high-quality imitations of these important traits, learners choose the best tutor in their population to imitate. In swamp sparrows, artificially slowed renditions of natural songs (<40% of the natural speed) are not imitated [39], suggesting that learners at least have a minimum set of criteria for tutor song choice. Moreover, in captivity, young males in this species given only accelerated tutor songs will attempt to imitate higher trill rates than they are capable of producing, resulting in atypical syntax with a few trilled notes punctuated by longer silences [40]. Although such “broken” songs are not encountered in nature, young sparrows are able to imitate them in the lab, suggesting both a flexibility that could enable novel song adoption, and a directional preference for copying faster trills. In another species of sparrow, artificial tutor songs with a single note removed were much less likely to be imitated than complete tutor songs, suggesting another kind of threshold for selecting high-quality songs to imitate [41]. In addition to judging their songs, juvenile birds might also gauge potential tutors using social cues [42,43]. This is somewhat analogous to cultural processes of prestige bias and success bias in humans [44,45]: in addition to judging tool quality (*z*), humans can also select tutors based on social factors, without invalidating Henrich’s model of cumulative cultural change.

When we apply Henrich’s adaptation of the Price equation to birdsong using a generic metric of song quality (*z*), we find, as expected, that the mean skill level of the population decreases over time when population sizes are small (lower *N*), or when traits are more difficult to imitate (higher α), and the mean skill level increases when population sizes are larger or when traits are easier to imitate (Figure 2). The tradeoff between population size (*N*) and trait difficulty (α) is shown by the white boundary in the heatmap, where Δ*z* = 0 and the skill level remains constant. Above this boundary (Δ*z*>0), the trait is expected to decline in quality over time. Conversely, increasing variance in learning (β) tends to promote an increase in *z* over time for a given population size (Figure S1).

**Figure 2.**
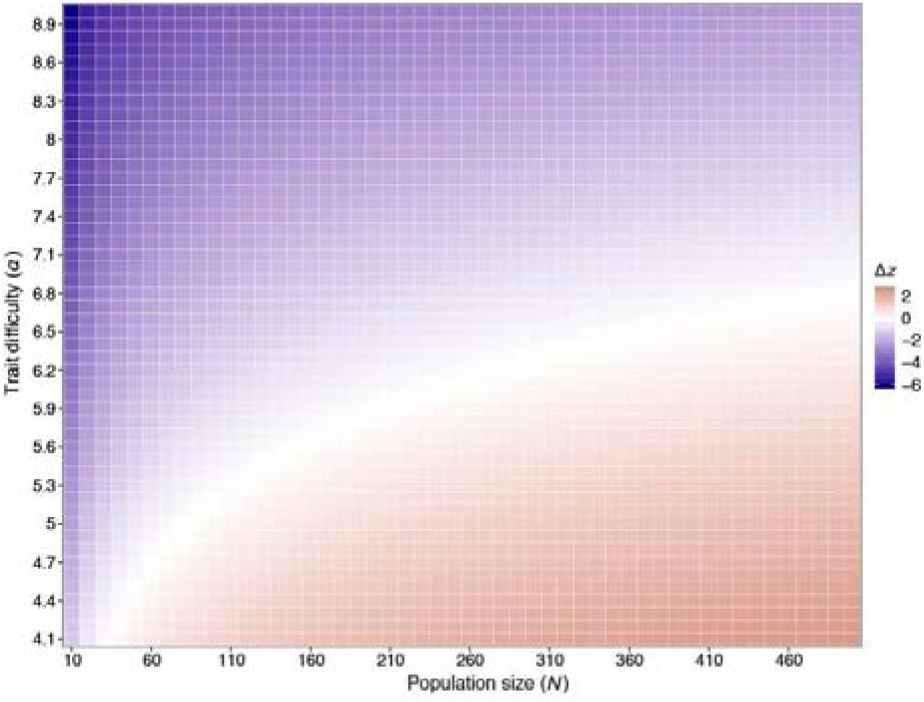
The effect of population size (*N*) and trait difficulty (α) on the change in mean trait skill level per generation (Δ*z*), where β=1. When population size is small, the mean change in learned skill is negative each generation (blue), even for lower values of α (easier traits). As population size increases, cultural traits that are increasingly difficult to copy (higher α) can be maintained and improved (red, positiveΔ*z*).

### Application to specific birdsong characteristics

So far, *z* has referred to any generic song trait that is under selection. Next, we apply a variation of this model to specific aspects of learned song. One aspect of birdsong that lends itself well to the Price equation model of cultural evolution is trill rate. This component of song is constrained by morphological limits [46], but also by the quality of the tutor song available: swamp sparrows presented with only artificially slowed songs could not produce songs reaching natural performance levels, although they did improve on their “tutor’s” performance [39]. Moreover, trill rate is functionally important for mate attraction [28,30,35] and territory defense [47–49]. However, several bird species show geographic variation in the rate or presence of trills [50–52], raising the question of why these seemingly important features of song are present (or more elaborated) in some populations and not others.

In Figure 3A, we show 20 stochastic replicates of possible trait trajectories over 60 generations in a population of *N*=100 individuals, starting from a uniform population trill rate of 4 notes per second (*z*=4). A population of size 100 is realistic and not computationally prohibitive, and the values of α=5 and β=1 represent a “boundary” at which minor parameter changes lead to changes in trait value (see Figures 2, S1). In the majority of replicates, the mean *z* of the population decreased over 60 generations; the mean of all replicates was a trill rate of 2.8 notes/second. This result suggests that, at these cognitive parameters (α and β), a population greater than 100 individuals is required to maintain trill rate at or close to the physiological limit. It is important to note that for most learned birdsong traits, physical or physiological limits exist that will constrain song expression; e.g. bill size determines how rapidly a male can trill [53,54], while neurophysiology places limits on auditory sensitivity [55,56]. Regardless of this upper limit, in our model, smaller populations experience a decline in trill rate as learners, over successive generations, largely fail to exceed the skill level of the best tutor of their generation. Thus, over time trills may disappear altogether, as the rate becomes so low as to no longer function in mate attraction. Indeed, one replicate in this simulation reached a mean *z*<0 before 60 generations (red line), which we represent as the loss of the trait. While the precise values are in some sense arbitrary (100 birds may be adequate to maintain most learned vocal behaviors in nature), this example illustrates the general principle that smaller populations are more likely to lose difficult-to-imitate traits than large ones.

**Figure 3.**
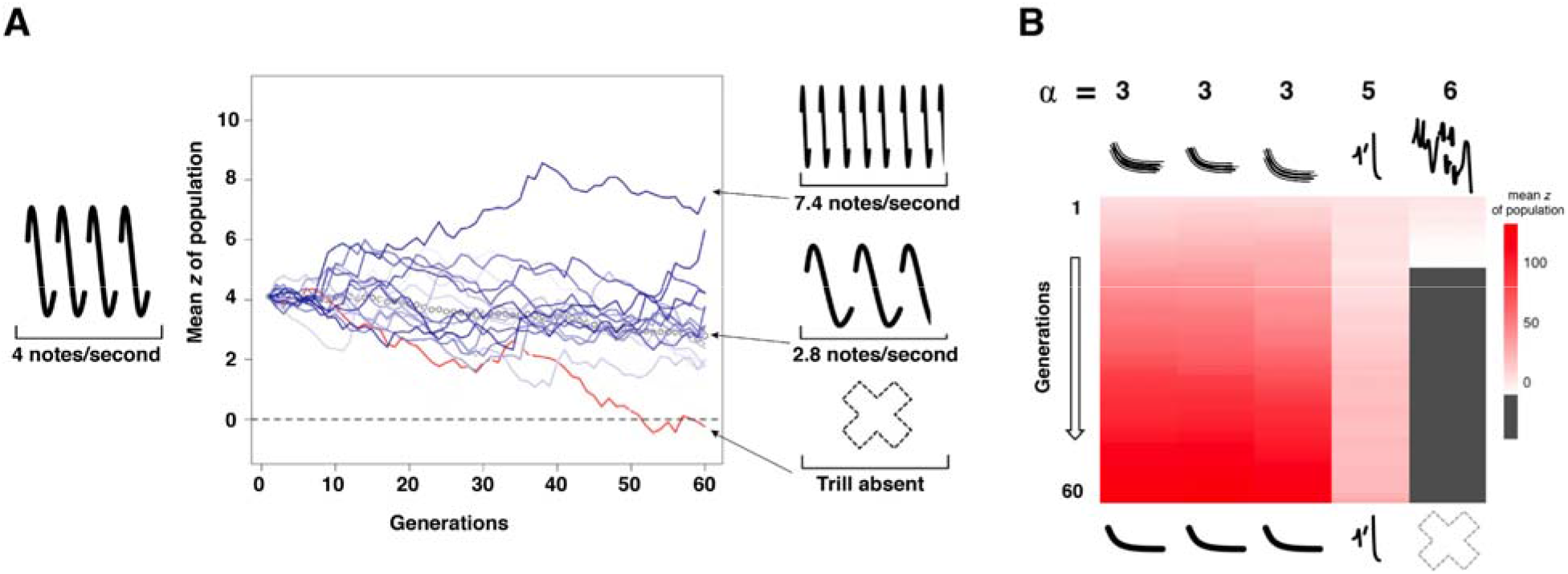
**(A)** The term *z* (a measure of skill level in a culturally transmitted trait) in the Price equation here represents the trill rate (number of notes repeated per second). Lines show the mean *z* of each replicate population over time. Open circles represent the mean *z* value across a**ll** 20 replicates for that generation. The range of final *z* values (maximum, mean, and minimum) after 60 generations are noted on the right y-axis. For all replicates, α=5, β=1, and *N*=100. (**B)** An ancestral song composed of 5 syllables of varying difficulty (□) is learned by a population **of** 100 individuals over 60 generations. The mean population value for *z* for each syllable is indicated by color over time, with red indicating higher values of *z*. In the context of birdsong evolution, *z*≤0 can be thought of as a syllable disappearing from a population’s song repertoire.

Even for song characteristics that are more difficult to quantify than trill rate, this model makes useful predictions about the direction of song evolution over time. For example, we can envision a bird’s song composed of multiple syllables that vary in their difficulty to learn. In the language of birdsong literature, this would be a more syntactically complex song than one composed of a single repeated note. Like trill rate, syllable and song complexity has been shown to play a role in mate attraction for some species, although the strength and ubiquity of this effect is contested (see meta-analyses by [10,57]). For example, female Bengalese finches prefer songs with higher complexity, but stressful conditions early in life limit a male’s syntactical complexity, indicating that these songs are more difficult to produce (large α) [58]. Repertoire size is another attractive feature of song that could be negatively affected by small population size. Although repertoire size *per se* may not be a culturally transmitted trait, the availability of diverse models to copy necessarily limits the achieved repertoire size of imitative learners, as seen in marsh wrens given a small pool of tutors from which to learn [14].

In the event of demographic changes such as population bottlenecks, our model predicts that syllables with a high trait difficulty (large α) would be the most likely to disappear from the population, while those that are simpler to learn (smaller α) would be maintained even after population size shrank. This differs from the prediction of song component loss through neutral processes (akin to genetic drift), because the loss of syllable types is not random. Rather, we would expect that more difficult-to-imitate syllables would be disproportionately lost if population size is reduced. This is true even if producing these syllables is beneficial to the singer. We illustrate one such scenario in Figure 3B, with a hypothetical birdsong initially composed of three different syllable types, each with its own imitation difficulty (α value). We show the results in a single population over 60 generations, in which each note is learned independently following the Price equation. Over 60 generations, the terminal syllable (which is the most difficult to imitate) rapidly decreases in *z*, while the middle syllable remains mostly the same and the initial notes increase in *z* value (Figure 3B). Note that the exact interpretation of t**he** *z* value in this context is more complex than in the case of the trill. One way to conceptualize a syllable with a larger *z* value is that the syllable is more salient or attractive to receivers (for example, one that is faster, more stereotyped, or more complex). In contrast, a syllable with a lower *z* value may be less salient to receivers (and perhaps disappears entirely from the song when *z=*0).

This model shows that a population of 100 is more likely to maintain song components below some threshold degree of difficulty (here, the threshold is 3<α<6).

## Song learning in a social network

Henrich’s application of the Price equation necessarily simplifies the nature of social interaction in a population. A more sophisticated model would include an estimation of the size of both the overall population and the number of individuals accessible for copying at each generation. Kobayashi and Aoki [59] devised such a model, in which individuals are randomly assigned *k* individuals out of the total population as potential cultural tutors, and choose the best tutor (highest *z*) from among those *k* individuals. Below we extend the methods of [59] in two ways to apply social network methodology to the question of how song might evolve in socially structured populations.

### Disentangling population size (N) from connectivity of a network (degree)

A key characteristic of social networks is degree, or the number of connections possessed by each member (node) of the network. (Networks with different degrees are illustrated in Figure S2). To tease apart the impact of local social network dynamics from overall population size on song evolution, we modeled two different network scenarios. First, we simulated a network with *N*=100 in which every individual was connected to twenty five others in the population (degree of 25). Initially, all individuals are assigned *z*=0. Thus, in the first time step of our model, individuals are assigned an identical tutor randomly from the 25 connected individuals. In all subsequent time steps, each individual selects the individual with the highest skill level (*z*) of its 25 connections to be its tutor. The likelihood that a pupil exceeds the tutor’s *z* value is defined by α and β, as before.

Second, we compared the above network to a fully connected network with *N*=26, such that each individual still has 25 connections, but they are to every other individual in the population. We found that the larger network with a population of 100 and degree of 25 consistently achieves a higher mean *z* value after 50 generations than a network of 26 fully connected individuals (Figure 4a), showing that there is indeed a “benefit” to being a part of a larger population, even if individuals in both populations only have 25 individuals to learn from at each time step. This benefit, as expected, decreases as the subset *k* approaches the size of the whole population (Figure 4b).

**Figure 4.**
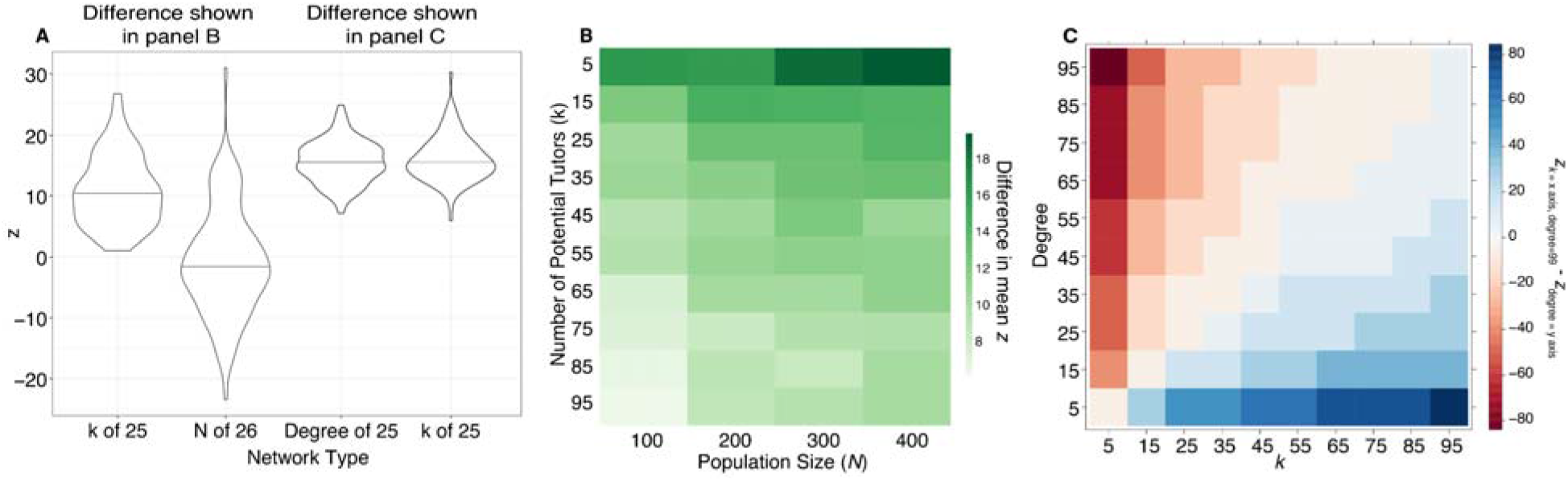
**(A)** Violin plots show the mean *z* in networks based on 100 simulations run for 50 generations. From left to right, the plots show the means of: a100-member network with a uniform degree of 25; a (fully connected) 26 member network with a uniform degree of 25; a 100-member network with 25 static connections; and a 100-member network where each node forms 25 connections randomly each generation. **(B)** The difference between the final mean *z* of a population after 50 generations with the total N and *k* on the x- and y-axes respectively, and a population with the same parameters but a total N equal to *k*. Values shown are the mean of 20 replicates. (**C**) The difference in mean *z* after 50 generations for populations where *N*=100, between networks with a *k* between 5 and 95 (where *k* potential tutors are sampled randomly from the network each generation), and networks with degree between 5 and 95 (where the same connections are maintained for all 50 generations).

### The effect of randomized versus consistent tutors

Next, we investigated two different scenarios in which only a subset of individuals in the network were available as potential tutors. In the first, we created a 100-member network and ran a simulation exactly as before, with each individual connected to 25 others (i.e. the network had a uniform degree of 25). Again as before, these individual connections were retained for the entire run (here, 100 generations). At each generation, each individual chooses the highest *k* among its connected nodes, and based on that maximum value, generates its new *z* value (“learns”) according to the Price equation. We replicated this simulation 100 times for 100 generations each.

We also created 100-member networks where each individual had the opportunity to learn from 25 individuals randomly drawn from the population in each generation (*k*=25). In contrast to the previous scenario, the 25 individuals were picked randomly for each individual during each learning event. We found that a fully connected network in which *k* individuals are randomly selected as potential tutors, and a network with a degree of *k*, yield the same mean *z* after 100 generations, as illustrated in Figure 4c. This suggests that whether the pool of tutors is static over multiple generations, or shuffled each generation, does not affect the overall outcome of song evolution. In other words, re-sampling tutors for each new generation of individuals does not provide any benefit for trait evolution by increasing the effective population size. Figure 4d illustrates that this pattern holds for values of *k* and degree between 5 and 95; when *k* and degree are the same, after 50 generations the difference in mean *<u>z</u>* is close to zero (light-colored diagonal line).

### The relationship between trait quality and difficulty: dynamic α as a function of *z*_max_

Following Henrich [15], we have so far assumed that the value of α for a given trait remains constant over time, regardless of the average skill level or complexity of that trait in the population. However, it is plausible that if the skill level of a trait decreases (for example, trill rate decreases or syllables become less structurally complex), learners will find it easier to successfully imitate. In this case, we would predict a positive correlation between α and *z*: as the quality of the trait increases (higher *z* values), the trait difficulty should also increase (higher α) and vice versa. Below, we model this scenario by modifying α at each generation, according to how far above or below the initial z value the current maximum z is at that generation (Figure 5).

**Figure 5.**
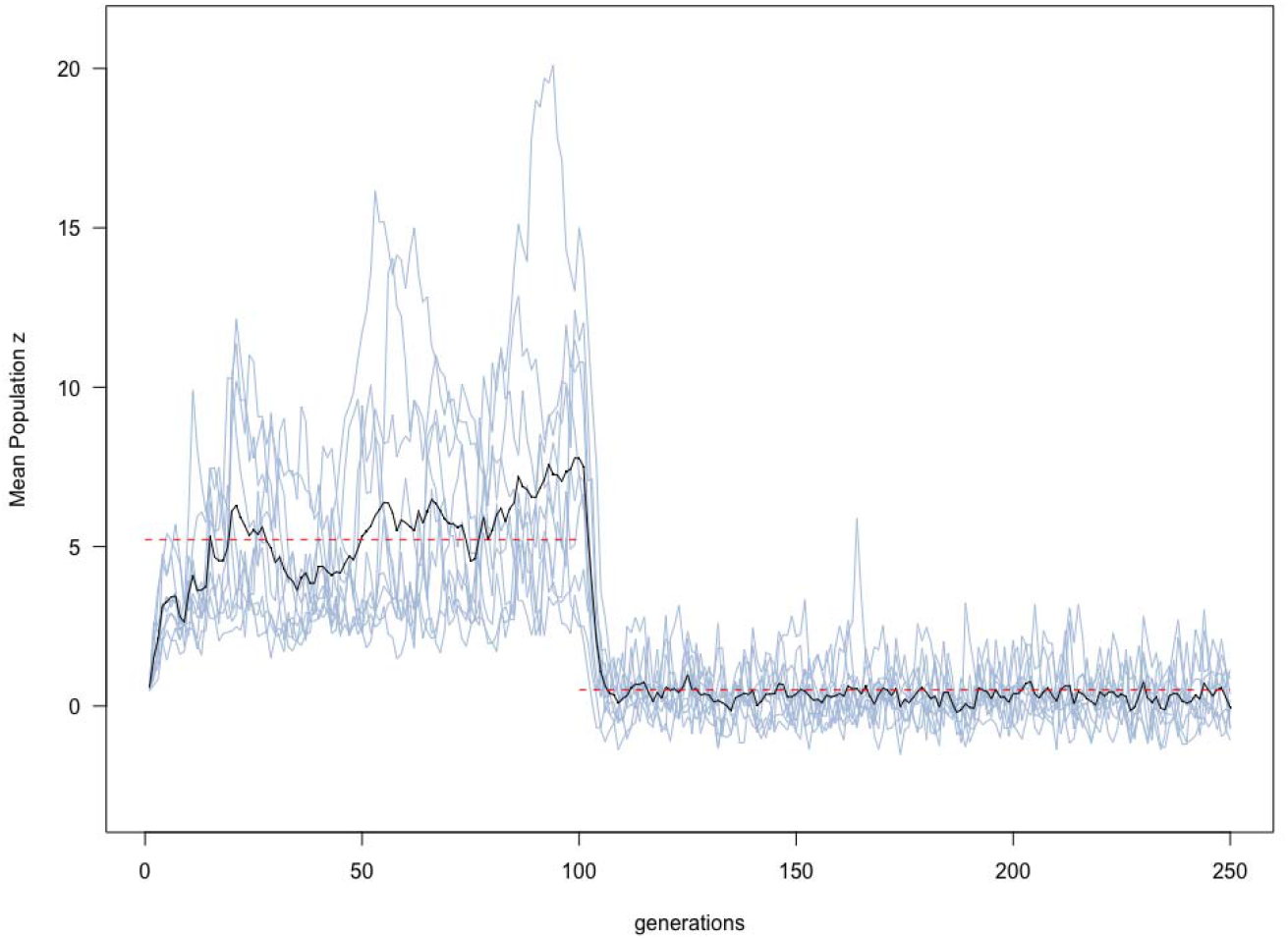
In these ten simulations, α varies between 3 and 7, changing each generation depending on the mean value of *z*, β=1, and the population shrinks from 500 to 50 at generation 50. Change from the original *z*_max_ (10) is used as an index to increase or decrease within a range of possible values of α. Red dotted lines indicate the mean *z* for the pre- and post-bottleneck populations.

Incorporating this plausible correlation between *z* and α produced two culturally and biologically relevant patterns. First, when α increases with *z*, mean trait performance improves until reaching a plateau, rather than a biologically unrealistic linear increase *ad infinitum* as seen in Henrich’s original model. As the quality of a trait improves in a population, the trait becomes progressively more difficult to learn, acting as a check on the unbounded increase in trait quality seen in the original model. Secondly, simplified traits may be maintained after drastic decreases in population size, rather than being lost altogether. In other words, as trait quality or performance declines in a small population, the trait becomes progressively easier to transmit and is rescued from loss. This concords with the relatively stable song traditions observed in most oscines, even in small or isolated populations.

## Discussion

Analogies between research fields can be useful in sparking the application of novel techniques to longstanding problems and encouraging interdisciplinary thinking. It is thus worth periodically revisiting the scientific analogies we use most, and adopting new and complementary ways of thinking about well-studied traits. Birdsong has often been compared to human language [60,61] and music [62], both symbolic traits; these analogies have no doubt encouraged the many decades of birdsong research in animal behavior, neuroscience, and psychology. We argue here that reframing birdsong as a functional tool is likely to spur further fruitful research. We draw an analogy between birdsong and human technologies—both of which are fitness-enhancing, culturally transmitted traits—and leverage models from cultural evolutionary theory to shed new light on the evolution of animal behavior. We have shown, using a relatively simple model based on the Price equation, that the size and network stability of a population can influence the retention of song traits over time. Population size and network structure in turn interact with cognitive parameters, such as the difficulty of copying a certain song feature and the variance of copying attempts, to shape song evolution. The interaction of demographic and cognitive parameters, we argue, is likely to affect the evolution of signals in oscine songbirds in similar ways to their effects on the evolution of human cultural traits: the structure of a population and the properties of its learned behaviors both play a role in cultural evolutionary dynamics. Specifically, populations that are small or sparsely connected should tend to lose their most difficult-to-copy traits over time, even if these traits continue to be advantageous to their bearers.

Empirical support for the hypothesis that population size affects song evolution can be found in studies of island populations of songbirds. For example, after a chain of island colonizations, chaffinches showed simpler phrases and trills than mainland populations [63]. Studies of this kind are hard to replicate, as they rely on large-scale phylogeographic events, but other song studies conducted on islands have shown rapid changes in songs [64,65] and in the associated recognition behavior [66]. Some common patterns emerge; for example, in a comparison of 49 pairs of mainland and island species, island species were less likely to sing rattles or trills [67], which are fast, complex sounds and thus potentially the most difficult to learn and produce. There are already hints that difficult-to-produce elements (such as trilled notes) may be vulnerable to loss in small wild populations not restricted to islands, as predicted by our model; e.g. golden-crowned sparrows in the furthest northwest breeding population lack terminal trills or buzzes in their songs [51]. Similarly, in the barn swallow, the populations that have the shortest and least complex songs also have the slowest trills [50]. In species of conservation concern, a common pattern observed when songbirds experience population bottlenecks is the shrinking of repertoire size [65,68,69]. In one study, after successive translocations of the North Island saddleback (*tĩeke*), high-pitched calls became lower pitched before eventually disappearing ([65], K. Parker, personal communication); it remains unknown whether these lower-pitched calls are easier to produce. One complication in detecting this pattern in more common species is that apparently small, isolated populations may in fact be in contact with a larger number of potential tutors throughout their lifetime, for example during migration or wintering [70]. Determining the demographic history, as well as the current effective number of tutors, of populations that have slow or no trills, versus those with rapid trills, could provide a test of the hypothesis that trills are lost when population size falls below a certain threshold.

A key assumption of Henrich’s model is that trait difficulty (α) stays constant for a given trait over time; in reality, a trait may become easier to learn as it simplifies. We modelled this variation and found an intuitive and important result: *z* approaches a minimum value as it becomes easier to replicate, enabling the trait to be stably maintained, though in a simpler form, in small populations that might have lost the trait when α was constant (i.e. the population maintains a small but positive *z* instead of a negative *z*). However, even when trait difficulty varies with trait quality, the trait can be lost from the population (*z* approaches a stable, but negative, value) under some parameter combinations. That simpler renditions of a species-typical song could be copied and maintained is consistent with empirical work in swamp sparrows [40], which shows that “poor” copies of song (with broken syntax, to accomodate trill rates beyond performance limits) are accepted as models by young sparrows and retained in future generations.

While population size has often been invoked in human cultural evolutionary theory as a factor in the maintenance or loss of learned skills, the existence of a straightforward relationship between population size and cultural complexity in humans has been contested [15,16,24,71–75]. Empirical research in humans has provided evidence both for and against such a relationship (see [73] for a review); similarly, there is also debate around the relationship between population size and maintenance of a large inventory of sounds in language [76,77]. The effects of population size may be complicated by other properties of a population, such as migration rates and connectivity. For example, a recent study showed that multiple, partially connected human groups can maintain a higher diversity of solutions to a complex problem [78]. This framework offers a promising perspective for investigating the origin and maintenance of dialects in many bird species, where spatial heterogeneity in song types can persist in the absence of obvious physical barriers.

We cannot rule out that there is cultural selection directly on elements of song, as in some aspects of human language [79], although there is no reason to suppose this will tend towards simplifying songs. Very rapid changes in song cultures do occur (e.g. [70]), and in the absence of obvious anthropogenic factors, such changes may be tied to cultural selection. Anthropogenic changes to the soundscape may also be a major factor in selection on song elements in the future, including elements that persist in the absence of noise [80] and may be transmitted socially [7]. Although our model does not address the effects of cultural selection or biased transmission on birdsong, clearly these mechanisms are extremely important for understanding evolution of learned vocalizations.

Bird species vary in the relative importance of song for mate attraction. Determining the relationship between song elaboration and female choice in the field, which would provide an key empirical test of the model we propose, is an important unresolved problem. A review of repertoire size and female choice studies in songbirds [27] highlights the discrepancy between laboratory studies of repertoire size, which often find a positive association between repertoire size and female preference, and field studies, which find such an association much more rarely. One explanation that these authors put forward is that there could exist “preferences that do not translate to choices” — that females in the field, while they may perceive differences in song repertoire, and even prefer larger repertoires, are also influenced by many other interacting factors, such as territory quality, in making their ultimate choice of mate [27]. We believe this principle likely applies to many aspects of female choice in the field. The strength of selection on song, and whether it arises from mate choice, intrasexual competition, or other mechanisms, likely affects how closely the assumptions of this model apply. Species with strong directional selection on song via mate choice by females, which most closely fits with the classic Fisherian runaway model, seem most likely to conform to the predictions we describe above. The stronger the link between song traits and fitness outcome, the more we might expect song to behave in a tool-like way. We encourage researchers interested in vocal learning and other forms of nonhuman cultural evolution to consider what novel predictions could be made in their system by embracing this subtle but important change of perspective.

