## Supplemental Methods and Figures 1 & 2 for "Ornament, armament, or toolkit? Modelling how population size drives the evolution of birdsong, a functional cultural trait"

**Supplemental Methods: Price Equation details**

In Henrich’s mathematical model, a learned trait (such as a netweaving technique, or the straightness of an arrow shaft, p. 200 of ref. [[16]](https://paperpile.com/c/4u2UeG/MnLHo)) gradually improves or declines across generations in a culture, depending on the interaction of three factors: 1) how difficult the trait is to accurately copy (α), 2) how widely learners vary in their attempts to copy the trait (β), and 3) how many learners and potential tutors exist in the current population (*N*), a non-cognitive factor that nevertheless plays an important role in cultural change. Henrich describes the relationship of these terms in a modified Price equation:

$\Delta\underline{z}= -\alpha+\beta$(ε + ln(*N*))	(1)

Here $\Delta\underline{z}$ is the population-level mean change in the skill level of a culturally transmitted behavioral trait, and ε is Euler’s constant ($\approx0.577)$. This equation represents a model of social learning where all individuals attempt to imitate a trait produced by the most competent individual in the previous generation, and there is no limit to the number of individuals that can imitate the same tutor. Population size *N* is important because larger populations have more individuals, increasing the probability that an individual produces a superior imitation. It is important to note that there is no direct inheritance in this model, genetic or otherwise; each pupil attempts to imitate the best tutor (*z_max_*) in the population, and in each generation the resulting *z* value for each pupil is drawn from a Gumbel distribution centered at *z_max_*$-\alpha$ (Figure 1).

When $\Delta\underline{z}$ > 0, the mean skill of the population increases, including the skill of the best tutor in the next generation (Figure 1B2). Conversely, $\Delta\underline{z}$ < 0 results in a decline in the population’s skill over time, shifting the distribution to the left.

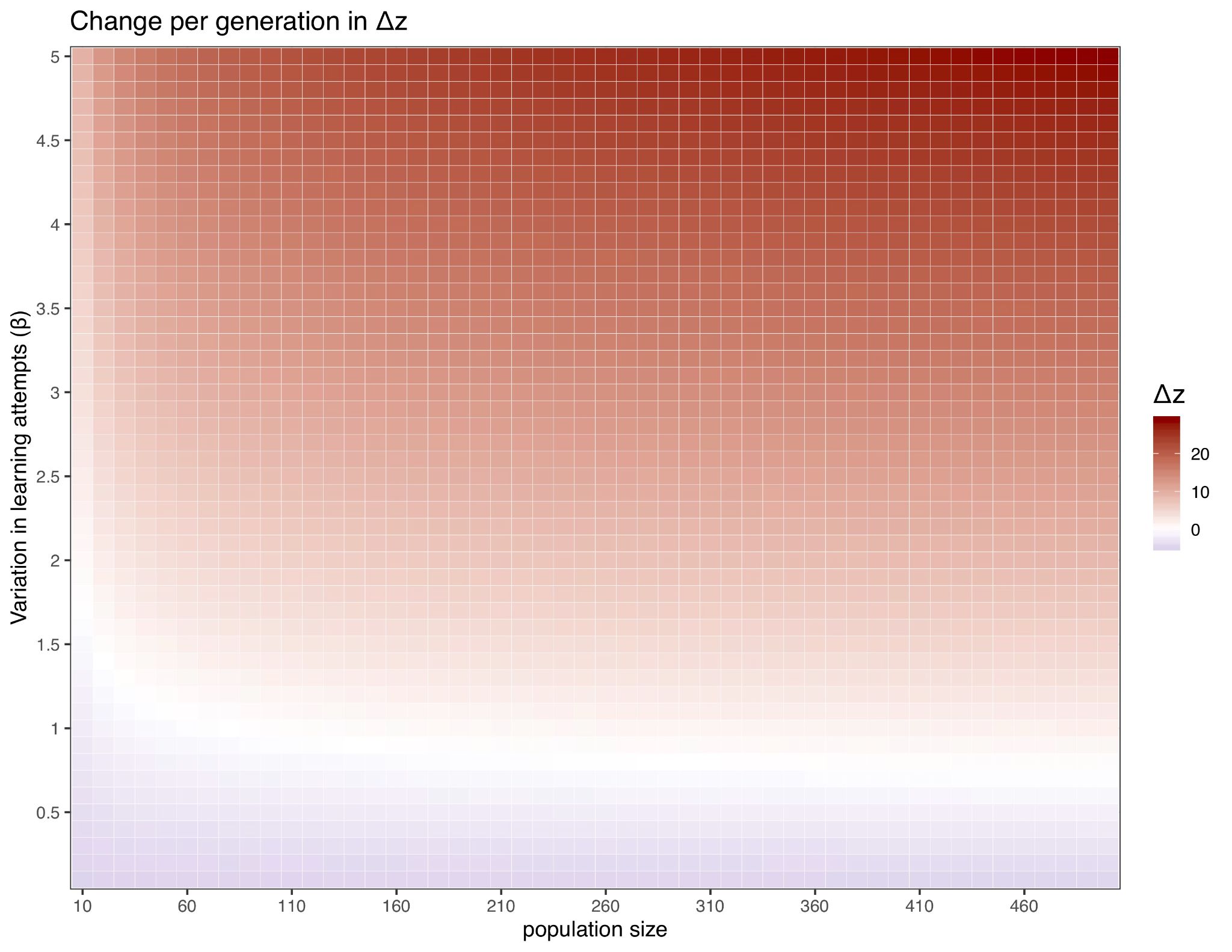

**Figure S1.** Increased learning variance (β) promotes the cumulative increase in trait values (*z*). Here, α = 5 and β varies between 0 and 5. As population size increases, cultural traits with lower learning variance (smaller β) can be maintained and improved (red, positive$\Delta z$).

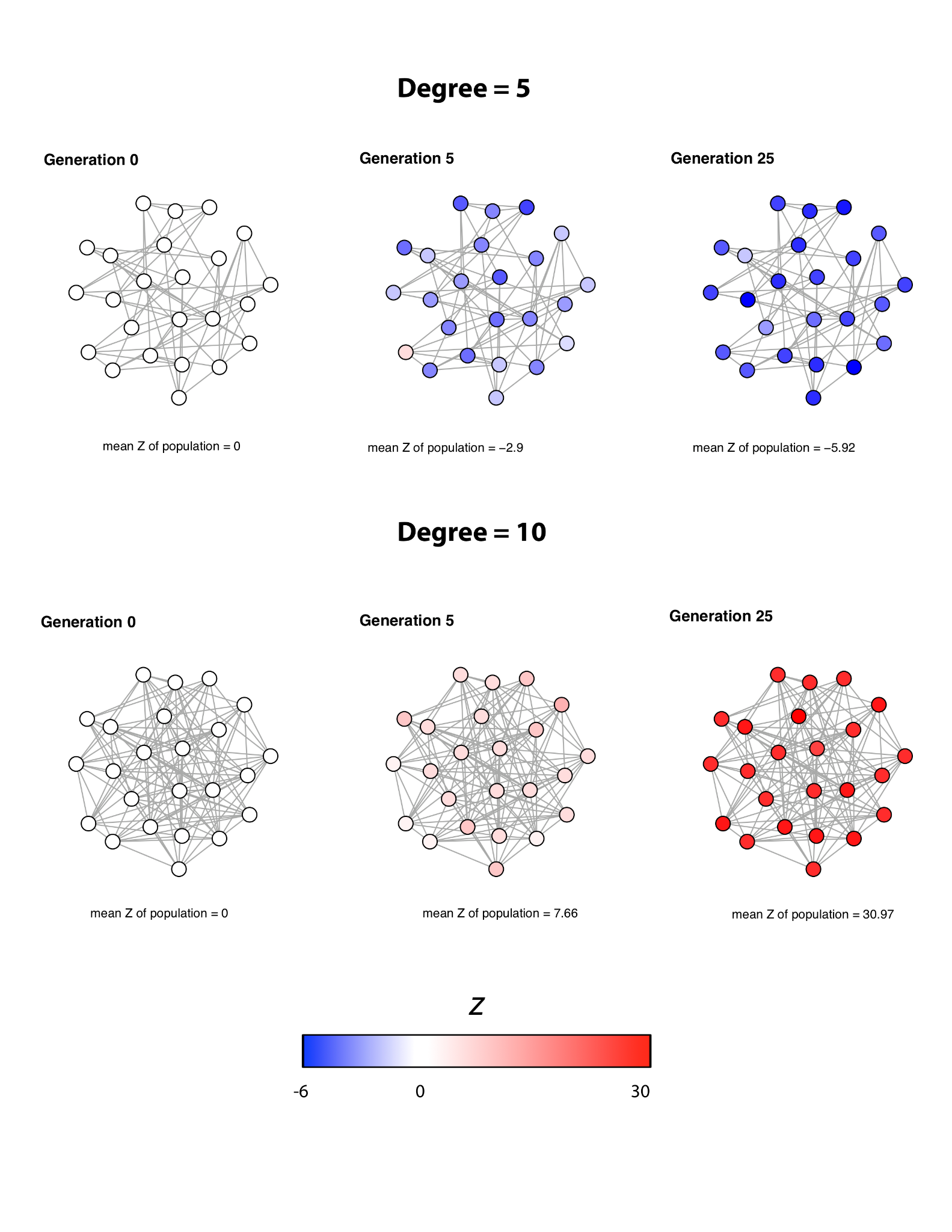

**Figure S2.** The simulated change in *z* in a population of 24 individuals where each member has a degree of 5 (top) or 10 (bottom). All individuals begin the simulation with a *z* of 0. The color of each node represents that individual’s *Z* value at a given timestep, and the mean *z* of the population is shown below each graph. Over 25 generations, the population with the smaller degree decreases in mean *z* (blue), while the larger degree network increases in mean *z* (red).
